# Multidimensional analysis of a social behavior identifies regression and phenotypic heterogeneity in a female mouse model for Rett syndrome

**DOI:** 10.1101/2023.06.05.543804

**Authors:** Michael Mykins, Benjamin Bridges, Angela Jo, Keerthi Krishnan

## Abstract

Regression is a key feature of neurodevelopmental disorders such as Autism Spectrum Disorder, Fragile X Syndrome and Rett syndrome (RTT). RTT is caused by mutations in the X-linked gene Methyl CpG-Binding Protein 2 (MECP2). It is characterized by an early period of typical development with subsequent regression of previously acquired motor and speech skills in girls. The syndromic phenotypes are individualistic and dynamic over time. Thus far, it has been difficult to capture these dynamics and syndromic heterogeneity in the preclinical *Mecp2*-heterozygous female mouse model (Het). The emergence of computational neuroethology tools allow for robust analysis of complex and dynamic behaviors to model endophenotypes in pre-clinical models. Towards this first step, we utilized DeepLabCut, a marker-less pose estimation software to quantify trajectory kinematics, and multidimensional analysis to characterize behavioral heterogeneity in Het over trials in the previously benchmarked, ethologically relevant social cognition task of pup retrieval. We report the identification of two distinct phenotypes of adult Het: Het that display a delay in efficiency in early days and then improve over days like wild-type mice, and Het that regress and perform worse in later days. Furthermore, regression is dependent on age, behavioral context, and is identifiable in early days of retrieval. Together, the novel identification of two populations of Het suggest differential effects on neural circuitry and opens new directions of exploration to investigate the underlying molecular and cellular mechanisms, and better design experimental therapeutics.

## Introduction

Regression is defined as a loss of previously acquired motor skills over time and is a behavioral hallmark of neurodevelopmental disorders (Charman et al., 2002; Goin-Kochel et al., 2014; McVicar & Shinnar, 2004; Thurm et al., 2018). Despite decades of clinical observations that regression is prevalent in many neurodevelopmental disorders and that the occurrence of regression is higher than previously reported, the underlying cause of regression in most neurodevelopmental disorders remains elusive (Goin-Kochel et al., 2014; Goldberg et al., 2008; Ozonoff et al., 2018; Ozonoff & Iosif, 2019). Genetic mutations in proteins important for transcriptional regulation and experience-dependent synaptic plasticity in the brain are associated with higher risk for regression (Goin-Kochel et al., 2017; Tammimies, 2019). Amongst these targets is the X-linked gene Methyl-CpG-Binding-Protein 2 (MECP2), the monogenic cause for Rett syndrome (RTT), and a “hotspot” gene associated with other disorders such as Severe Neonatal-Onset Encephalopathy, PPM-X syndrome and microcephaly (Amir et al., 1999; Gonzales & LaSalle, 2010; Lambert et al., 2016; Rett, 1966; Schüle et al., 2008).

Most surviving RTT patients are girls and women who are heterozygous for *MECP2* mutations (Kirby et al., 2010). Female patients typically survive into middle age, and exhibit sensory processing, cognitive and motor deficits throughout life (Buchanan et al., 2019; Djukic et al., 2012; Djukic & Valicenti McDermott, 2012; Key et al., 2019; LeBlanc et al., 2015; Merbler et al., 2020; Neul et al., 2010; Nomura, 2005; Peters et al., 2015; Stallworth et al., 2019; Symons et al., 2019). In these females, random X-chromosome inactivation leads to mosaic wild-type MECP2 expression and consequently, a syndromic phenotype. Broadly, Rett syndrome is diagnosed from two types of clinical presentations: 1) girls present with a short period of typical development, followed by regression, and expression of stereotypic sensory, motor, speech and cognitive impairments; or 2) developmental delay and expression of stereotypic sensory, motor, speech and cognitive impairments (Charman et al., 2002; Cosentino et al., 2019; Djukic & Valicenti McDermott, 2012; Einspieler & Marschik, 2019; Hagberg et al., 1983; Han et al., 2012; Neul et al., 2010, 2023; Symons et al., 2019). Regression typically occurs between 6-18 months of age, and as late as 4 – 8 years from individual case studies (Buchanan et al., 2019; Han et al., 2012). These prolonged timelines of vulnerability suggest that the increased cognitive load required during early development while mastering motor control through exploration and interacting with the social environment might reveal specific features over ages. Though RTT is diagnosed in early development, regression in specific skills or learning is also observed throughout life. Currently, the underlying neural pathways that manifest in developmental delay, or regression after typical development and throughout different phases of life are unknown.

Preclinical *Mecp2* rodent models recapitulate many of the phenotypic features of RTT such as sensory processing, social communication, and motor deficits (Durand et al., 2012; Goffin et al., 2012; Ito-Ishida et al., 2015; Krishnan et al., 2017; Lau, Krishnan, et al., 2020; Lee et al., 2017; Lo et al., 2016; Mykins et al., 2023; Orefice et al., 2016; Stevenson et al., 2021; Su et al., 2015; Veeraragavan et al., 2016; Xu et al., 2022). Thus far, three other published papers have shown possible regression phenotypes, for breathing abnormalities in a mouse conditional knockout model, learned forepaw skill involving seed opening in a female rat model, and in skilled motor learning in female mice (Achilly, Wang, et al., 2021; Huang et al., 2016; Veeraragavan et al., 2016). In these models, deterioration in learned skills was noted over multiple weeks. However, reliably identifying regression within a social behavioral task in apt preclinical models, with shorter time resolution in order to measure impact of therapeutics, has remained a challenge.

Towards this end, we have established an ethologically relevant social behavioral assay called pup retrieval task as a model to study cellular and neural circuitry basis for social cognition, complex sensorimotor integration and experience-dependent plasticity (Dvorkin & Shea, 2022; Krishnan et al., 2015, 2017; Lau, Krishnan, et al., 2020; Lau, Layo, et al., 2020; Mykins et al., 2023; Stevenson et al., 2021; Xie et al., 2023). During this task, female mice integrate sensory cues to execute goal-directed motor sequences to retrieve scattered pups back to their home nest (Alsina-Llanes et al., 2015; Beach & Jaynes, 1956; Champagne et al., 2007; Champagne & Curley, 2016; Dunlap et al., 2020; Hemel, 1973; Koch & Ehret, 1989; Lonstein et al., 2015; Marlin et al., 2015; Stern, 1996; Stern & Mackinnon, 1978). Using end-point metrics such as latency index and errors during retrieval, we found that 6 week-old adolescent *Mecp2*-heterozygous female mice (Het) were as efficient at pup retrieval as wild-type littermate controls (WT), and 12 week-old adult Het were inefficient at retrieval (Krishnan et al., 2017; Mykins et al., 2023; Stevenson et al., 2021). However, movement sequences during social behaviors are dynamic over time, with distinct behavioral sequences that are often ignored due to their inherent complexity (Stevenson et al., 2021).

The emergence of computational neuroethology tools allows for fast, reliable, and systematic detection of recorded movement over time to extract multiple metrics in different contexts. This extraction is critical for identifying unique sub-populations and assessing the therapeutic value of manipulations for pre-clinical animal studies for psychiatric disorders (Luxem et al., 2023; Popovitz et al., 2021; Shemesh & Chen, 2023; Tanas et al., 2022; Yamamoto et al., 2018). Thus, to capture the complexity of individual variations in a mosaic *Mecp2*-heterozygous population over multiple days of pup retrieval behavior, we used DeepLabCut, a marker-less pose estimation software, to derive animal trajectories, and performed multidimensional analysis of trajectory kinematics in adolescent and adult female WT and Het (Mathis et al., 2018; Nath et al., 2019). The goal of this study was to determine if DeepLabCut can provide better behavioral characterization and aid in identifying regression in this heterogeneous population of Het.

Here, we report the identification of robust phenotypic variation and regression in genotypic adult Het. Particularly, we identified genotypic Het that had milder phenotypes, similar to adult WT in many features (Het non-regressors, Het-NR), and another distinct group of genotypic adult Het that exhibited severe inefficiency in consolidating the pup retrieval movements over days (Het regressors, Het-R). Interestingly, the regression was dependent on context as we observed WT, Het-NR and Het-R clustered together in Principal Component space during habituation and isolation phases, suggesting Het-R have specific issues likely in integrating sensory and motor sequences during the decision/execution phase of retrieval. Additionally, this inefficiency in consolidation was specific to adulthood as adolescent Het behaved similarly to adolescent WT, indicating genotypic Het are able to perform this complex pup retrieval task at an earlier time point in life, and are not able to maintain goal-directed movement skills as they age. The novel identification of two populations, Het-NR and Het-R, suggests differential effects on neural circuitry and opens new directions of exploration on cellular mechanisms for compensation and maladaptive plasticity. Together, these results emphasize the need to analyze dynamics in individual variations and context-dependency while considering therapeutic and treatment options.

## Materials and Methods

### Animals

We used the following mouse strains: CBA/CaJ (JAX:000654) and *Mecp2^heterozygous^* (Het) (C57BL/6J, B6.129P2©-Mecp2^tm1.1Bird^/J, JAX:003890) and wild-type littermate controls (WT) (Guy et al., 2001). All animals were group-housed by sex after weaning, raised on a 12/12-hour light/dark cycle (lights on at 7 a.m.) and received food and water ad libitum. Behavioral experiments were performed using 6 week-old (adolescent) and 10-12 week-old (adult) Het and WT between the hours of 9 a.m. and 6 p.m. (during the light cycle). All procedures were conducted in accordance with the National Institutes of Health’s Guide for the Care and Use of Laboratory Animals and approved by the Institutional Animal Care and Use Committee at the University of Tennessee-Knoxville.

### Pup Retrieval Task

#### Behavioral Paradigm

The pup retrieval task was performed as previously described (Mykins et al., 2023; Stevenson et al., 2021). Two 6-week-old or 10-12 week-old naïve female littermates (one WT and one Het) with no prior experience of pup retrieval were co-housed with a pregnant CBA/CaJ female (10-12 weeks of age) 3-5 days before birth. Once the pups were born (Postnatal Day 0; D0), we performed the pup retrieval task once a day for six consecutive days, in the home cage which was placed inside of a sound- and light-proof box. The behavioral task was performed as follows (Figure 1a.): one mouse was placed in the home cage with 3-5 pups (Habituation; 5 minutes), the pups were removed from the nest (Isolation; 2 minutes), and the pups were scattered by placing them in the corners and center, allowing the mouse to retrieve pups back to the nest (Retrieval; 5 minutes at max). The nest was left empty if there were fewer than five pups. The assay was performed in the dark and recorded using an infrared camera (Foscam Wired IP Camera) during all phases of each trial. If all pups were not retrieved to the nest within 5 minutes, we removed the mouse and placed pups back in the nest in preparation for the next mouse. While one of the females was performing the pup retrieval task, the mother, her pups, and the other adult were housed in a separate new cage. After the behavior was completed, all mice and pups were returned to the home cage

**Figure 1.**
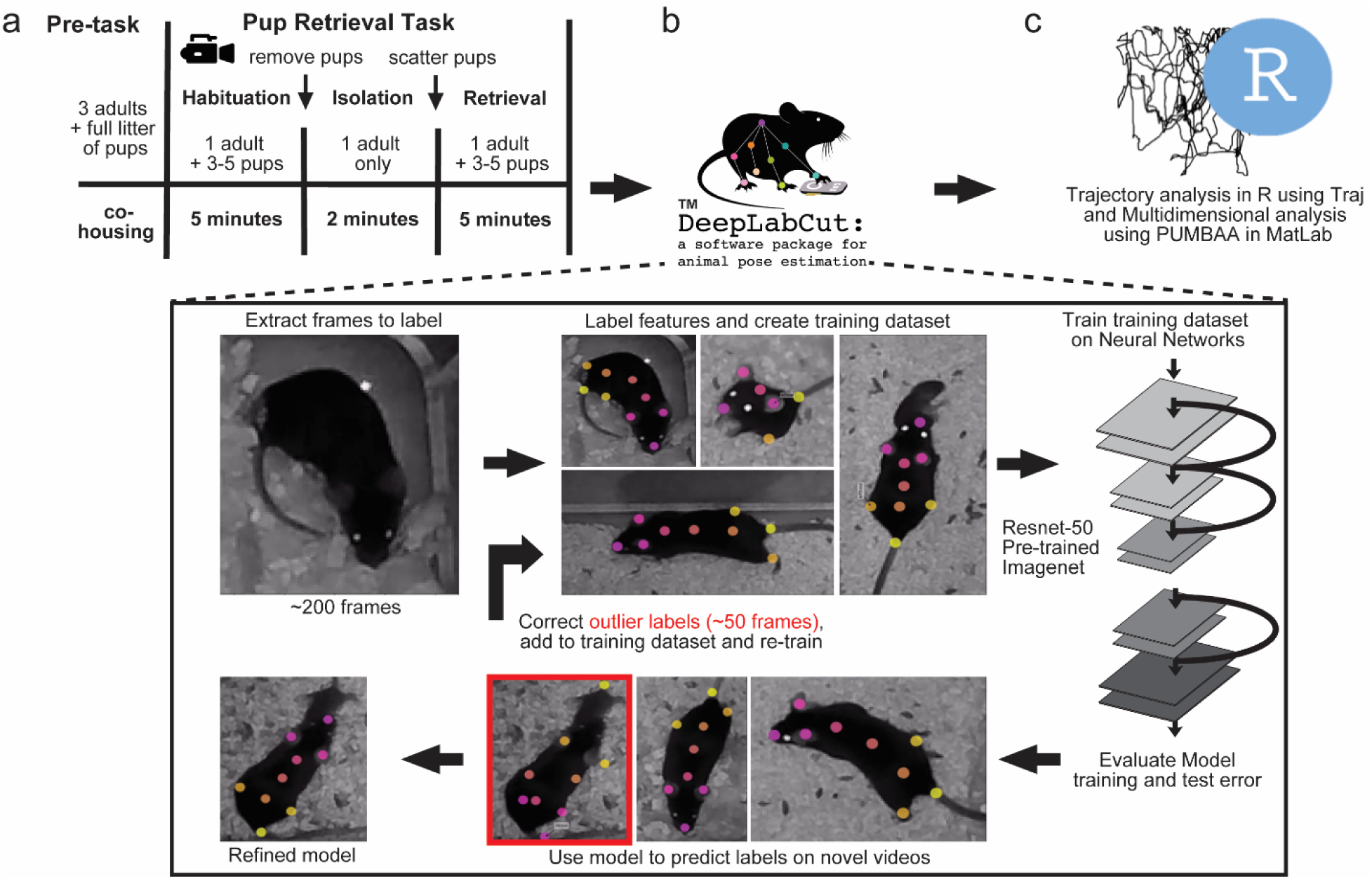
DLC derived pipeline for multidimensional analysis of trajectories. a) Schema depicting behavior setup. We performed and recorded habituation, isolation and pup retrieval in 6 weeks old (WO) adolescent and 10-12 WO adult female WT and *Mecp2^heterozygous^* (Het) mice for 6 consecutive days (see methods for details). b-c) We trained videos of behavior in single animal DeepLabCut (DLC) (b: top) to predict pose of the animal. b-bottom: features of interest were labeled and trained in DLC to build pose models which were then refined with more training to correct for errors. From our models we generate pose estimation and extract features of interest such as the nose to analyze trajectories of our mice in R using the Traj package (*McLean and Skowron Volponi 2018*) (c) and perform multidimensional behavioral analysis adapted from Tanas et al. 2022.

#### Behavior Analysis

All recorded videos were coded so the analyzers were blind to the identity of the mice. Each video was manually scored using a latency index (the amount of time to retrieve all pups back to the nest out of 5 minutes) and the number of errors that represent adult-pup interactions that did not result in successful retrieval).

Latency index was calculated as: *Latency index = [(t1 – t0) + (t2 – t0) + … + (tn - t0)] / (n x L)*

Where n = number of pups outside the nest, t0 = start time of trial (s), *tn* = time of the nth pup’s successful retrieval to the nest (s), L = total trial length (300 s). Statistical analyses and figures were generated using R studio.

#### Pose estimation generated from DeepLabCut

For body part tracking, we used DeepLabCut (version 2.2.0.3) (Figure 1b) (Mathis et al., 2018; Nath et al., 2019). We defined features of interest as the mouse’s nose, left ear, right ear, shoulder, spine 1, spine 2, left hind limb, right hind limb, and tail base. We labeled 200 frames of features of interest taken from 25 videos of WT and Het during pup retrieval phase and then 95% was used for training. We used a ResNet-50-based neural network with default parameters and trained for 720,000 iterations on Google Colab (Insafutdinov et al., 2016). We performed outlier correction to correct inaccurate predictions from the DeepLabCut model. In the Graphical User Interface (GUI), we refined 50 frames of predicted labels by moving the predicted label’s location to the actual position of the body part of interest (Figure 1b). We then retrained, validated with 1 number of shuffles, and found the test error of was 1.67 pixels, and train error was 1.66 pixels (image size was 640 X 480 pixels). We then used a p-cutoff of 0.9 to condition the X and Y coordinates for future analysis. This model network was then used to analyze novel behavioral videos of adult animals. We then repeated this process for 6-weeks-old adolescent pup retrieval behavior. We labeled 200 frames taken from 20 videos of 6-weeks-old pup retrieval behavior, trained for 800,000 iterations, refined 75 frames of predicted labels, retrained for 550,00 iterations, validated with 1 number of shuffles, and repeated this process to achieve test error of was 1.89 pixels, and train error was 1.82 pixels.

#### Pose estimation derived trajectory analysis

To quantify behavioral trajectories during habituation, isolation and retrieval, we used the Traj R package (McLean & Skowron Volponi, 2018) (Figure 1c). We filtered the X and Y coordinates of the nose from the DLC pose and quantified 13 Trajectory metrics (11 metrics were used for habituation and isolation as they did not have latency index and error) (Table 1). For habituation and isolation phases, we analyzed trajectories during the entire 5-minute and 2-minute trial phases, respectively. For pup retrieval phase, we analyzed trajectories during the act of retrieval; we stopped analyzing trajectories after all pups have been retrieved.

**Table 1.** Trajectory metric definitions.

| Metrics: | Definition: |
| --- | --- |
| Latency index | - The amount of time to retrieve all pups back to the nest out of 5 min (retrieval phase of behavior only, excluded in habituation and isolation) |
| Errors | - Number of physical interactions with pup during the retrieval phase which do not result in retrieval, or dropping of the pup during retrieval (retrieval phase of behavior only, excluded in habituation and isolation) |
| Distance | - Total distance covered (in meters) during phase of behavior. |
| Duration | - The total amount of time it takes to retrieve all pups, or the max 5 minutes (retrieval phase)<br>- For habituation, 5 minutes total<br>- For isolation, 2 minutes total |
| Mean speed | - Change in distance over time (scalar quantity) |
| Mean acceleration | - Acceleration- change in speed over time (scalar quantity) |
| Mean step length | - The average distance between two steps |
| Sinuosity | - Tortuosity of a random search path (Benhamou, 2004; Bovet & Benhamou, 1988)<br>- A function of both the mean cosine of turning angles and step length (Cheung et al., 2007; McLean & Skowron Volponi, 2018)<br>- Varies between 0 (straight) and 1 (random motion) |
| Straightness | - Minimum distance to get from point a to b (D) divided by displacement from origin to final destination (L) (Batschelet, 1981)<br>- Closer to 1, the more efficient the trajectory. The closer to zero, the more random the walk (Batschelet, 1981; Benhamou, 2004) |
| E<sub>max</sub> | - Dimensionless value that denotes the maximum expected displacement of a random path as a function of the number of steps (Cheung et al., 2007; McLean & Skowron Volponi, 2018)<br>- Values closer to 0 are more sinuous trajectories, larger values (approaching infinity) are straighter trajectories (Cheung et al., 2007) |
| E<sub>maxb</sub> | - The product of E<sub>max</sub> times the defined step size (Cheung et al., 2007)<br>- larger values (approaching infinity) are straighter |
| Nonlinearity | - Measure of directional change<br>- Angular change (in degrees) between two steps, divided by the time difference between the two steps (Benhamou, 2004; Kitamura & Imafuku, 2015) |
| Irregularity | - Standard deviation of directional change (nonlinearity) (Benhamou, 2004; Kitamura & Imafuku, 2015) |

### Multidimensional analysis of trajectories

For multidimensional analysis of behavior profiles, we performed data selection, standardization, principal component analysis (PCA), k-means clustering, and validation (Fig. 2b) (Popovitz et al., 2021; Tanas et al., 2022). We utilized PUMBAA (phenotyping using a multidimensional behavioral analysis algorithm, https://github.com/sidorovlab/PUMBA <u>A)</u> for all multidimensional analysis (Tanas et al., 2022).

**Figure 2.**
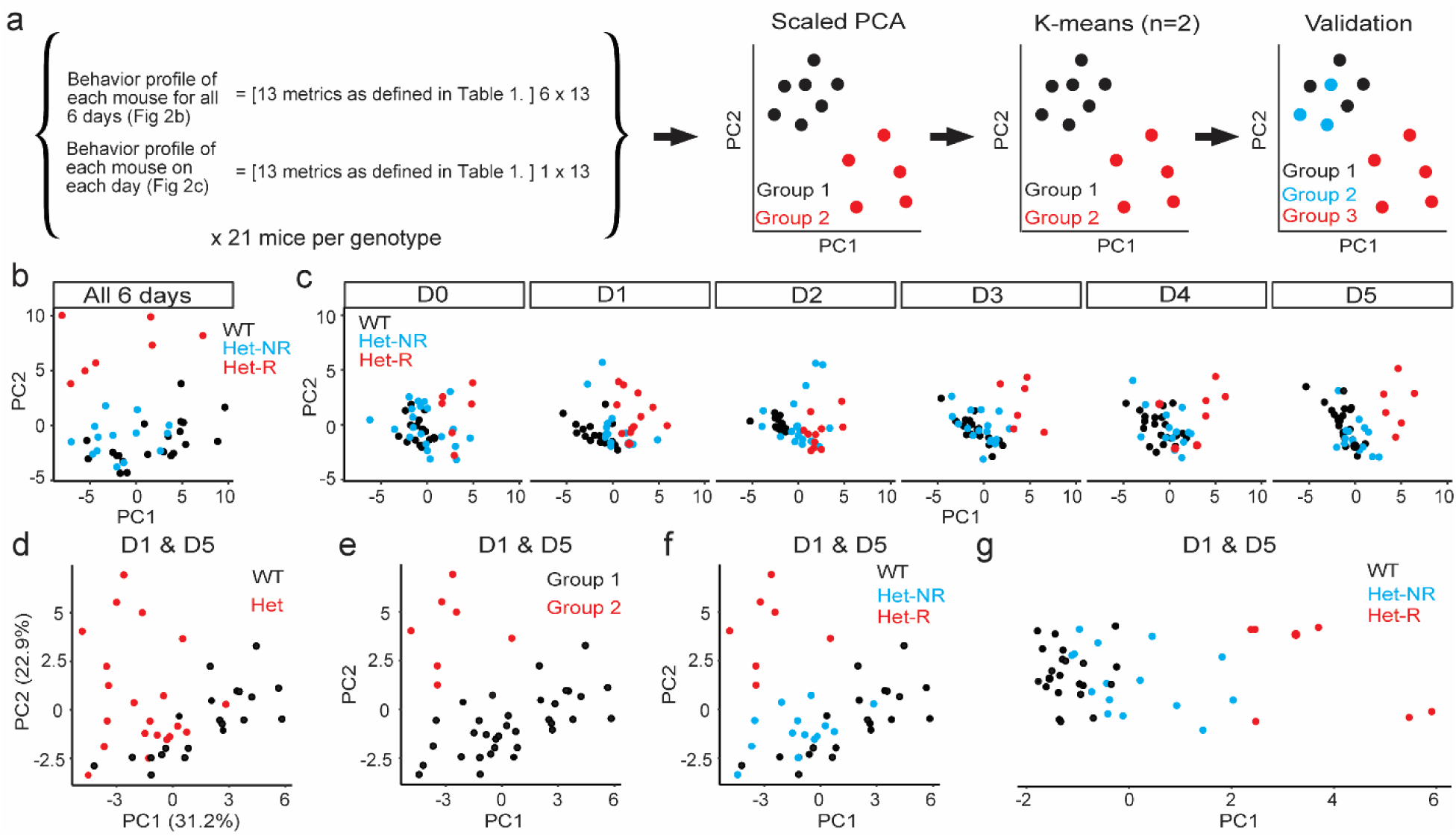
Multidimensional Analysis identifies two distinct phenotypes of Het. **a)** 13 trajectory metrics for all days of retrieval (78 total loadings) or 13 trajectory metrics on each day (13 total loadings) were standardized using z-scores for PCA (left). Example of analysis: each point represents one animal’s behavioral profile in PC space, colored by Group (Scaled PCA). Mice were clustered into two groups (Group 1= black, Group 2=red) using k-means to predict genotype (K-means). The predicted genotypes were validated by comparing animals to their genotype (WT=black, Het predicted correctly = red, Het predicted incorrectly = blue) (Validation). **b)** Multidimensional analysis for all days of retrieval revealed two groups of Het: Het that clustered with WT (Het-NR, blue) and Het that were distinctively different from WT and Het-NR (Het-R, red). **c)** PCA, k-means clustering, and validation for each individual day of retrieval (D0-D5). **d)** PCA of 13 trajectory metrics for D1 and D5 of retrieval (26 loadings) color coded by actual genotype (WT=black, Het=red). **e)** K-means clustering (*n*=2) of 13 trajectory metrics for D1 and D5 of retrieval (Group 1= black, Group 2=red). **f)** Validation of predictions supports two distinct groups of Het with reduced data complexity. **g)** Principal component 1 is sufficient to separate the two distinct groups of Het. For all panels: each dot represents an animal, color-coded by genotype and phenotype. WT (black, n=21), Het-NR (blue, n=14), and Het-R (red, n=7)

#### Data selection for retrieval

For individual and combined-days analyses of retrieval, 13 X number of days of metrics were included n multidimensional analysis or PCA analysis.

#### Data selection for habituation and isolation

For all days’ analysis, a total of 66 metrics (11 metrics across 6 days) were included in multidimensional analysis (Figure 4b, 4e). For combined D1 and D5 analysis, 22 metrics were included in multidimensional analysis (Figure 4c-f).

#### Standardization

All measures were standardized using z-score normalization (z = (data point − group mean) / standard deviation). This accounts for different unit magnitudes across metrics similar to previous studies (Popovitz et al., 2021; Tanas et al., 2022).

#### Principal component analysis and k-means clustering

We performed PCA using PUMBAA and extracted the amount of variance explained by each principal component (Figure 3b). We performed k-means clustering in principal component space using PUMBAA with k = 2 clusters in 2 Principal Component (2PC) space. To verify visual absence of clusters, we performed gap statistics test using the cluster package in R. Data visualization using ggplot were performed in R studio (Wickham, 2016).

**Figure 3.**
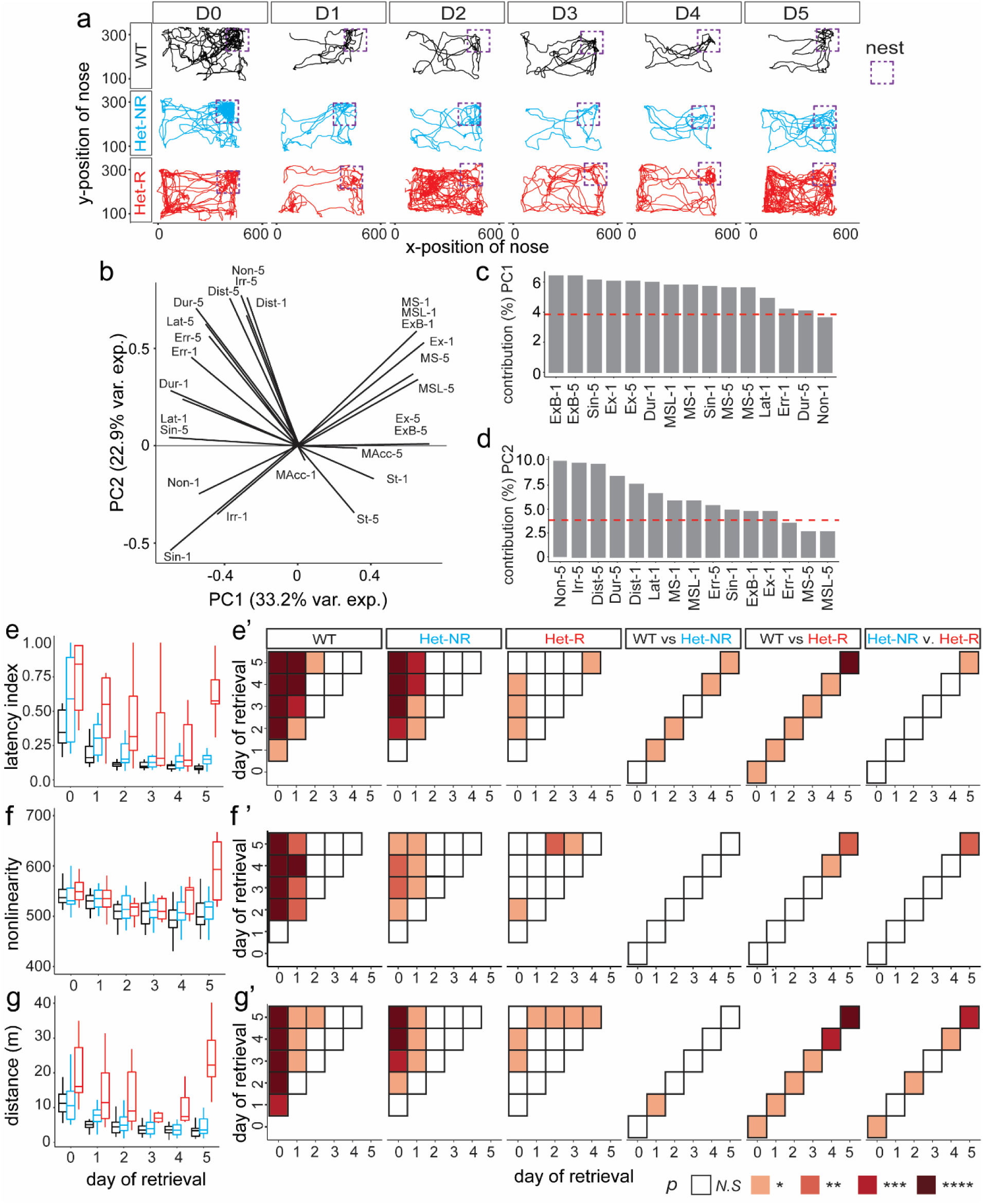
Multidimensional analysis distinguishes Het that improve and regress in behavioral efficiency over days. **a)** Example trajectories of WT (black), Het-NR (blue) and Het-R (red) x-y nose position during pup retrieval towards the nest (purple box) across 6 consecutive days of behavior. **b)** Variable correlation plots of trajectory metrics (see key below) from D1 and D5 represented in PC space. Variable vectors grouped in the same direction are positively correlated, and vectors in the opposite direction are negatively correlated. Variance explained (var. exp. %) represents the percent of variance explained by each PC component. **c-d)** Percent contribution of top 15 (out of 26) metrics to Principal Component 1 (c) and Principal Component 2 (d). The red dashed line represents the average expected contribution of each metric, everything above the red dashed line has considerable contribution to the component. **e)** latency index, **f)** Median distance nonlinearity and **g)** Median distance traveled during pup retrieval for each test day showed significant improvement in retrieval trajectories across test days for Het-NR. However, Het-R improved over days and regressed by D5 compared to D0 and were worse than WT and Het-NR. WT (black, n=21), Het-NR (blue, n=14), and Het-R (red, n=7). Kruskal-Wallis followed by uncorrected dunn’s test was performed to test for statistical significance between genotypes, and within genotype across days. Significance is plotted as correlation matrices colored coded by significance for **e’)** latency index, **f’)** nonlinearity and **g’)** distance. * p<0.05, ** p<0.01, *** p<0.001; **** p<0.0001, N.S. = not significant. b-d) Trajectory metric key: number following abbreviation indicates the day of retrieval. [Dist: Distance, Dur: Duration, Err: Error, Ex: Emax, ExB: EmaxB, Irr: Irregularity, Lat: Latency, MAcc: Mean acceleration, MS: Mean speed, MSL: Mean step length, Non: Nonlinearity, Sin: Sinuosity, St: Straightness]

#### Validation

We compared the genotypes to their predicted k-means cluster. We color-coded groups according to their predictions (WT predicted correctly= black, Het predicted correctly=red, and Het predicted incorrectly=light blue) (Figure 2). This allowed us to determine the accuracy of predicting genotypes in our model and revealed two distinct populations with the same genotype. Data visualization using ggplot was performed in R studio (Wickham, 2016).

#### Using predicted clusters to analyze behavior

Using the identified clusters of Het from Figure 2f, we performed trajectory analysis for metrics contributing significantly to the PCA. We conducted the Kruskal-Wallis test with uncorrected Dunn’s test to determine statistical significance of metrics of interest between WT and the two clusters of Het (Figure 3) on each day and within the same group across days. We used ggplot and the dunn.test package to perform statistical analysis and data visualization in R studio (Wickham, 2016).

## Results

### Robust identification of phenotypic heterogeneity in adult Het during pup retrieval

Previously, we reported that the adult female Het were inefficient at pup retrieval, as measured by time taken to retrieve pups back to the nest, and increased physical interactions between pups and adult that did not result in efficient retrieval (Krishnan et al., 2017; Stevenson et al., 2021). We also reported individual variations with possible regression phenotype over days in adult Het, from frame-by-frame manual analysis of goal-directed movements during pup retrieval in a small sample size (N=6 females per genotype) (Stevenson et al., 2021). These intriguing results identified the need for systematic, automated and unbiased analysis for large sample sizes over multiple days. Additionally, we sought to identify newer dynamic metrics in context-specific motor sequences, in addition to the established end-point measurements. To achieve these goals, here we used DeepLabCut to generate animal pose and quantified behavioral trajectory profiles across pup retrieval days in adolescent (N=9, WT and N=11, Het) and adult female WT and Het (N=21 per genotype) (Mathis et al., 2018) (Figure 1a,b). We incorporated animal trajectory analysis that included 13 metrics (Table 1), and adapted multidimensional analysis methods to cluster animal phenotypes, based on their genotypes and behavioral profiles (Figure 1c) (Benhamou, 2004, 2014; Bovet & Benhamou, 1988; Cheung et al., 2007; McLean & Skowron Volponi, 2018; Tanas et al., 2022). Together, we took 13 metrics from each day of retrieval (13 metrics per day) or across 6 days of pup retrieval (78 metrics total) to build a behavioral profile for each mouse (Figure 2a). We standardized all metrics using z-scores and performed multidimensional principal component analysis (PCA) (Figure 2a, Scaled PCA) (Popovitz et al., 2021; Tanas et al., 2022). We then used k-means to cluster mice into two groups based on genotype and validated clustering by comparing the predicted cluster group to the known genotype (WT vs Het) of the animal (Figure 2a, K-means and Validation, respectively). Based on this analysis, we identified a distinct group of Het clustered at the top of principal component (PC) space (Het-R, red) while another cluster with WT (Het-NR, blue) (Figure 2b). When we performed multidimensional analysis on each day of retrieval, we observed the Het-R cluster emerged as early as D1, however, it consolidated with WT and other Het on D2 and D3 and emerged again on D4 and D5 (Figure 2c). This suggests heterogeneity in phenotype is dynamic over days. Pairwise combinations between D0:D5, D1:D5, D2:D5, D3:D5 and D4:D5 (Figure supp. 1) indicated multidimensional analysis of D1 and D5 is sufficient to identify the two clusters of Het via PCA, k-means and validation (Figure 2d-f). Furthermore, PC1 alone distinguished the separation between the two clusters of Het (Figure 2g). Together, these novel results robustly quantify the known behavioral heterogeneity in adult Het over multiple days and identify regression in a subpopulation of adult Het.

### Identification of trajectory metrics that classify behavioral heterogeneity over days

In order to better characterize the three phenotypes (WT, Het-NR, Het-R), we plotted the trajectories, and observed striking differences in the movement patterns over days (Figure 3a). Similar to WT, Het-NR trajectories were random on D0 and consolidated to oriented and direct paths toward the nest by D5 (Figure 3a). Het-R also had random trajectories on D0 but showed instability in consolidating trajectories over time compared to WT and Het-NR (Figure 3a).

To determine the identity of the variables which contribute to the variance in PCA, we plotted the variable vectors in principal component (PC) space (Figure 3b) for D1 & D5, as this allowed us to isolate the regressors with the least amount of dimensions (Figure 2f). Sinuosity expected maximum displacement (emax) and expected maximum displacement B (emaxB) contributed to PC1, and distance, duration, nonlinearity, irregularity, latency index, and error contributed to PC2, whereas mean acceleration and straightness had negative correlations and did not contribute to either principal component (Figure 3b). Positive correlations between emax, emaxB, mean speed, mean step length, and negative correlation with sinuosity contributed to PC1 & PC2 (Figure 3c). Breakdown of contributions by PC1 (Figure 3c) and PC2 (Figure 3d) indicated nonlinearity, irregularity, distance, and latency contributed to the most variability in PC2.

We previously reported latency index, or the time it takes to retrieve all pups, as an endpoint to evaluate pup retrieval performance (Krishnan et al., 2017; Mykins et al., 2023; Stevenson et al., 2021). As a positive control, we also quantified latency index for this larger dataset. As expected, WT and Het-NR significantly decreased in latency index over days compared to baseline, here defined as D0 performance, within genotype (Figure 3e, e’, WT, Het-NR). This suggests learning and consolidation towards efficient pup retrieval task over days. However, Het-NR were significantly worse than WT on D1, D2, D4 and D5 (Figure 3e, e’, WT vs Het-NR), suggesting the possibility of intact learning and consolidation mechanisms, albeit not as well as WT controls. Similar to WT and Het-NR trends, Het-R latency index significantly decreased in early days (Figure 3e, e’, Het-R). However, Het-R latency increased from D4 to D5 and reverted to D0 levels. Het-R were significantly worse than WT on all days (Figure 3e, e’, WT vs Het-R) and were significantly worse than Het-NR on D5 (Figure 3e, e’, Het-NR vs Het-R). Together, these results suggest Het-R display delay in learning efficiency and are unable to consolidate efficient patterns over time.

Nonlinearity is a measure of directional change (Cheung et al., 2007; Kitamura & Imafuku, 2015; McLean & Skowron Volponi, 2018). A decrease in directional change is an indication that the path is more goal directed. WT and Het-NR significantly decreased in nonlinearity over days compared to baseline within genotype (Figure 3f, f’, WT, Het-NR), suggesting consolidation of more goal directed trajectories over days. There was no significant difference in nonlinearity between WT and Het-NR on all days (3f, f’, WT vs Het-NR). However, Het-R had significantly increased nonlinearity on D4 and D5 and were significantly worse on those days compared to WT and Het-NR (Figure 3f, f’, WT vs Het-R, Het-NR vs Het-R).

The total distance covered in a goal directed trajectory decreases with efficiency and is thus indicative of the efficiency of the path taken. WT and Het-NR significantly decreased their total distance covered over days, compared to baseline within genotype (Figure 3g, g’, WT, Het-NR), suggesting increased efficiency in determining the fastest path to retrieve pups back to the nest. Similar to WT and Het-NR, Het-R significantly decreased in total distance in early days compared to Het-R baseline (Figure 3 g, g’, Het-NR). However, Het-R significantly increased the total distance travelled on D5 compared to D1-D4, and was similar to total distance traveled at baseline (Figure 3 g, g’, Het-R). Het-R covered more total distance than WT on all days (Figure 3g, g’, WT vs Het-R) and were significantly worse than Het-NR on D0, D2, D4 and D5 (Figure 3g, g’, Het-NR vs Het-R).

Overall, compared to WT, Het-NR displayed mild and day-specific atypical metrics in latency index and distance travelled, while still improved from their own baseline performance in all metrics. However, Het-R had a larger latency index, made more directional changes, and covered more distance on each day of retrieval. The inability of Het-R to improve in latency (Figure 3e, e’, Het-R), nonlinearity (Figure 3f, f’, Het-R), and distance (Figure 3g, g’, Het-R) between D0 and D5, despite improved performance in intermediate days, indicates regression in this social motor task by D5.

### Features of Het regression are context-dependent

Given the dynamic changes in trajectory metrics during pup retrieval over days in the Het-R, we wondered if these features were specific to the act of pup retrieval, or other generalized motor features in this subpopulation. Thus, we performed principal component analysis of trajectories during the habituation and isolation phases of behavior (Figure 4). As latency and error metrics are specific to adult and pup interactions during retrieval, we excluded them in the multidimensional analysis. During habituation, the pups are grouped together in the nest and the adult can choose to interact/huddle with pups or not (Figure 1a). This serves as a control to determine if the context and action of retrieving pups distinguishes Het-NR and Het-R. We plotted the habituation trajectories (example plots for WT (black), Het-NR (blue) and Het-R (red)) and observed no differences in the patterns of their trajectories between D0 and D5 (Figure 4a). Both genotypes tend to move around the entire cage during the five-minute habituation phase. No distinct groups emerged between WT, Het-NR and Het-R from the differential PCA analyses (Figure 4b, c).

**Figure 4.**
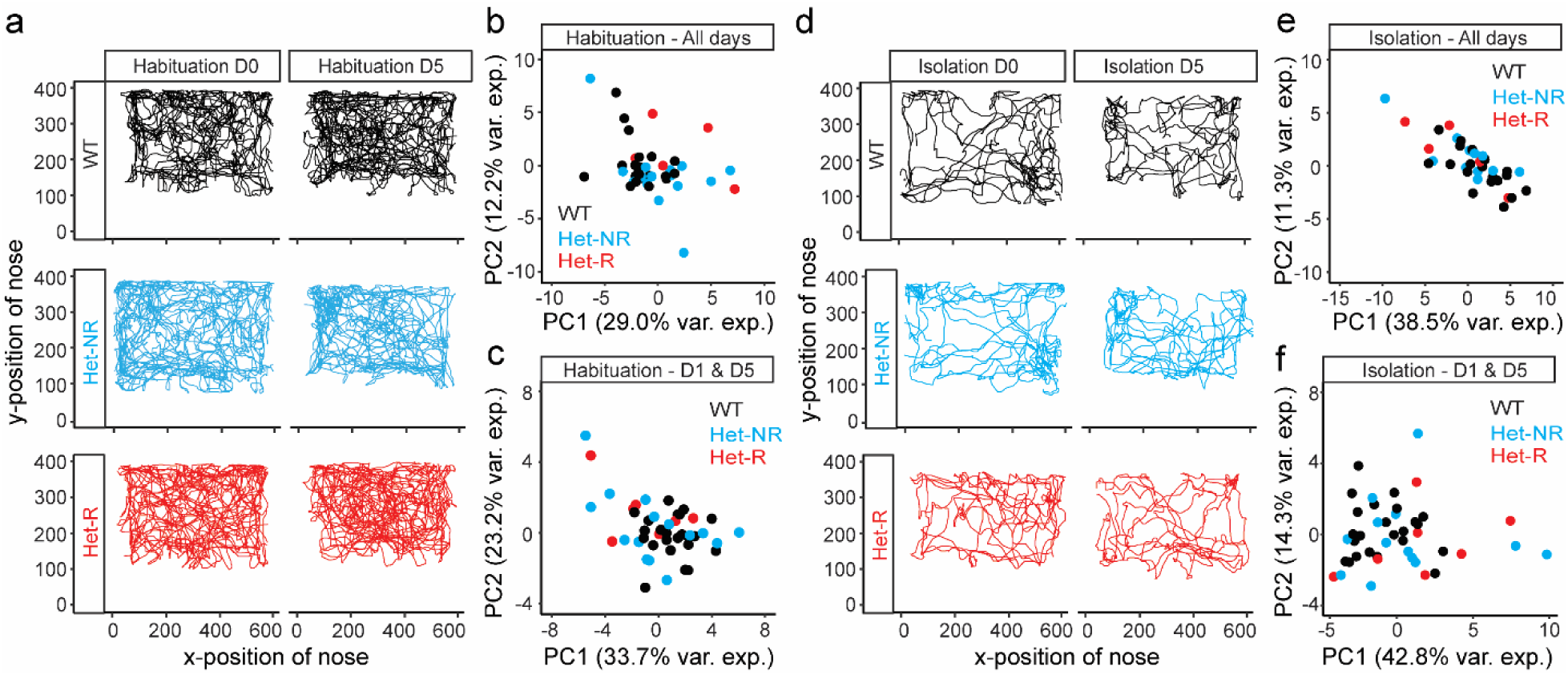
Regression is context-dependent. **a)** Example trajectories of WT (black), Het-NR (blue) and Het-R (red) x-y nose position during habituation on D0 and D5. **b-c)** PCA for all 6 days **(b)** and combined D1 and D5 **(c)** of habituation revealed no difference between identified groups during habituation (WT=black, Het-NR= light-blue, and Het-R=red). **d)** Example trajectories of WT (black), Het-NR (blue) and Het-R (red) x-y nose position during isolation for D0 and D5. **e-f)** PCA for all days **(e)** and combined D1 and D5 **(f)** of isolation revealed no difference between identified groups in the absence of pups. For **b-c, e-f**: each dot represents an animal, color-coded by genotype and phenotype. WT (black, n=21), Het-NR (blue, n=14), and Het-R (red, n=7)

During the isolation phase, the pups are removed from the home cage, and the adult is left alone for two minutes (Figure 1a). We plotted the isolation trajectories of the three groups and observed no differences in the patterns of their trajectories between D0 and D5 (Figure 4d). Similar to habituation phase, both genotypes moved about the cage with no pups present, and no distinct groups emerged from the isolation-specific PCA analyses (Figure 4e, f). These findings are consistent with the manual frame-by-frame analysis of maternal behaviors, which demonstrated WT and Het behave similarly during habituation and isolation (Stevenson et al., 2021). Together, these results suggest context-specific, atypical behavioral features of Het-R and Het-NR, and not generalized motor phenotype in genotypic adult Het. This is a critical finding which points to context-specific processing of sensorimotor information in genotypic Het and subsequent execution of specific motor sequences in a social context.

### Analysis of early days identifies individual variation in later adult Het regressors

In order to determine individual variations in Het, we plotted individual trajectories of Het-NR and Het-R over all days. As expected, we observed that Het-NR, such as animals 8, 20 and 21, displayed some variability in adapting over time, compared to the other Het-NR animals that plateaued and consolidated their efficiency in later days (Figure 5a). In contrast, we observed variability in individual trends in Het-NR. Animal 10 did not improve in pup retrieval efficiency over days, animals 3, 4, 6, 9, 19 improved over days but regressed by D5 and animal 2 improved over days but showed instability in D4 (Figure 5a’), suggestive of different neural mechanisms being disrupted in individual Het over days.

**Figure 5.**
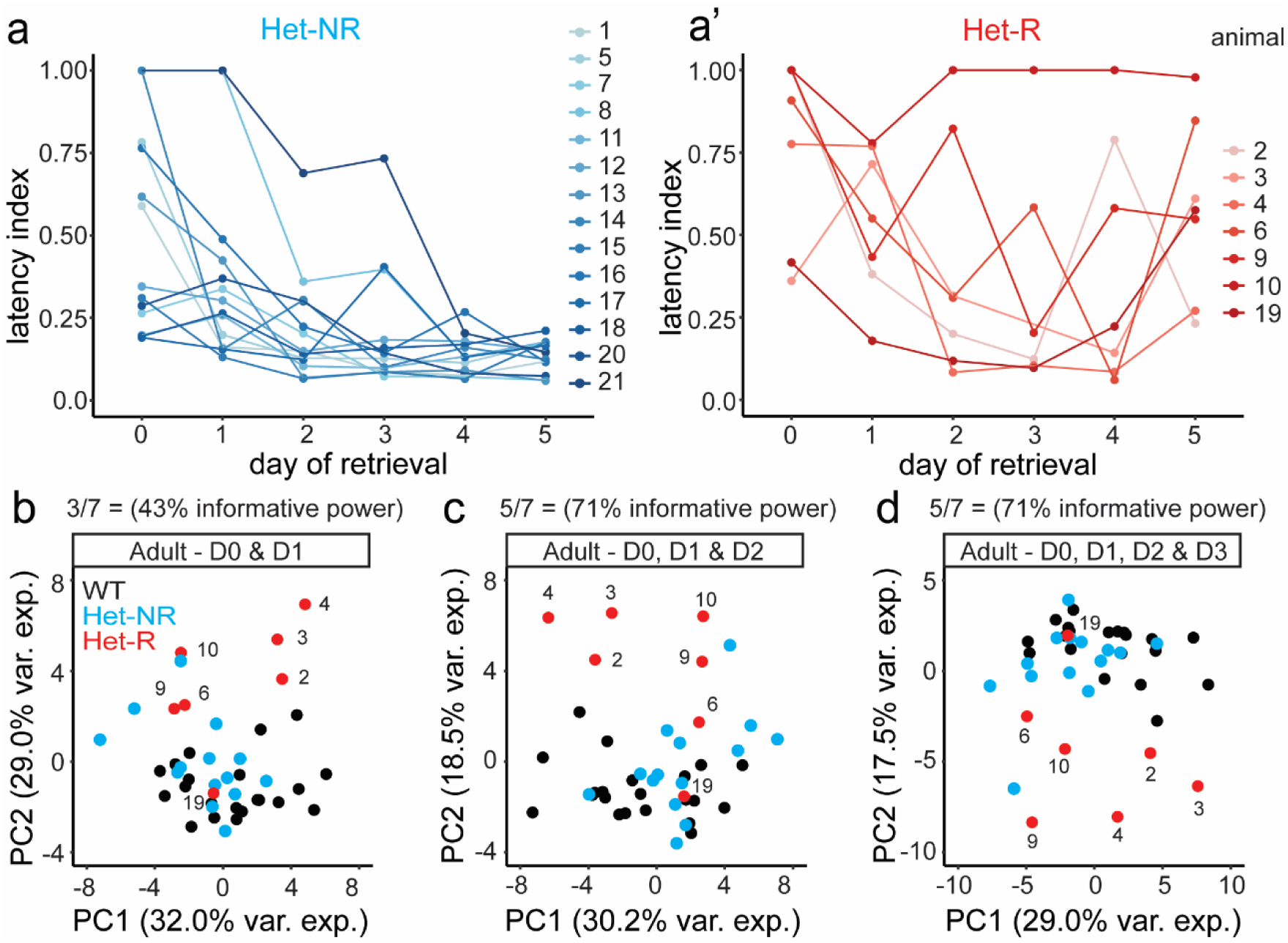
Performance in early days is able to identify future adult Het regressors. **a-a’)** Individual animal trend lines of latency index for the identified (a) Het Non-Regressors (Het-NR) and **(a’)** Het-Regressors (Het-R) from Figure 2 and 3. Trend lines are colored coded by animal number and highlight individual variation in latency index over 6 days for Het-NR (blue-gradient) and Het-R (red-gradient). **b-d)** PCA analysis for D0 and D1 (b), D0, D1 and D2 (c) and D0, D1, D2, and D3 **(d)** of retrieval. Analysis revealed that the first 3 **(c)** or4 **(d)** days of pup retrieval had a strong accuracy of 71% for identifying 5 out of 7 Het regressors that regressed on D5 of retrieval, while the first 2 days **(b)** had a lower accuracy of 43% (3 out of 7 regressors). Informative power was determined by counting the number of D5 regressors within each cluster and dividing by 7 (the total expected number of regressors). Dots represent each animal [WT (black, n=21), Het-NR (blue, n=14), and Het-R (red, n=7)], and Het-R are numbered to easily identify regressors over time.

While it has been difficult to phenotype regression in Het mouse models of RTT, it is an even greater challenge to determine the time, frequency, and duration to apply therapeutic intervention to mitigate regression before its onset. We thus asked if it was possible to identify the Het-R population (Figure 2; Figure 5a’) from the initial days of pup retrieval performance. Using D0 & D1 metrics (Figure 5b), we obtained weak informative power for identifying a majority of regressors, with 43% accuracy. However, combinations of D0-D2 (Figure 5c) and D0-D3 (Figure 5d) metrics elicited stronger informative power for identifying a majority of regressors early on, with 71% accuracy. Despite observations from select metrics that show Het-R trends like WT and Het-NR in early days (Figure 3e, f, g), regression in social behavior can still be identified using all metrics (Figure 5c, d). Together these results suggest that we can identify potential regressors with a smaller number of trials which has implications for designing preclinical therapeutic studies in the future.

### Experience dependent regression in Het is specific to age

We previously reported 6-week-old (adolescent) Het perform pup retrieval comparable to adolescent WT, using end-point metrics of latency index and error (Mykins et al., 2023). This is a particularly important result that shows that genotypic Het can perform as efficiently as the WT at an earlier age, which then results in regression features in adulthood. To determine if dynamic pose estimation metrics can identify early signs of regression in adolescent Het, we generated another deep learning model and pose for adolescent WT and Het animals, and quantified trajectory metrics. We did not observe any distinct genotype groups through PCA analysis (Figure 6a, b). In support of this finding, we performed the gap statistic test and determined that the optimal number of clusters between both genotypes was one (Figure 6 a’, b’).

**Figure 6.**
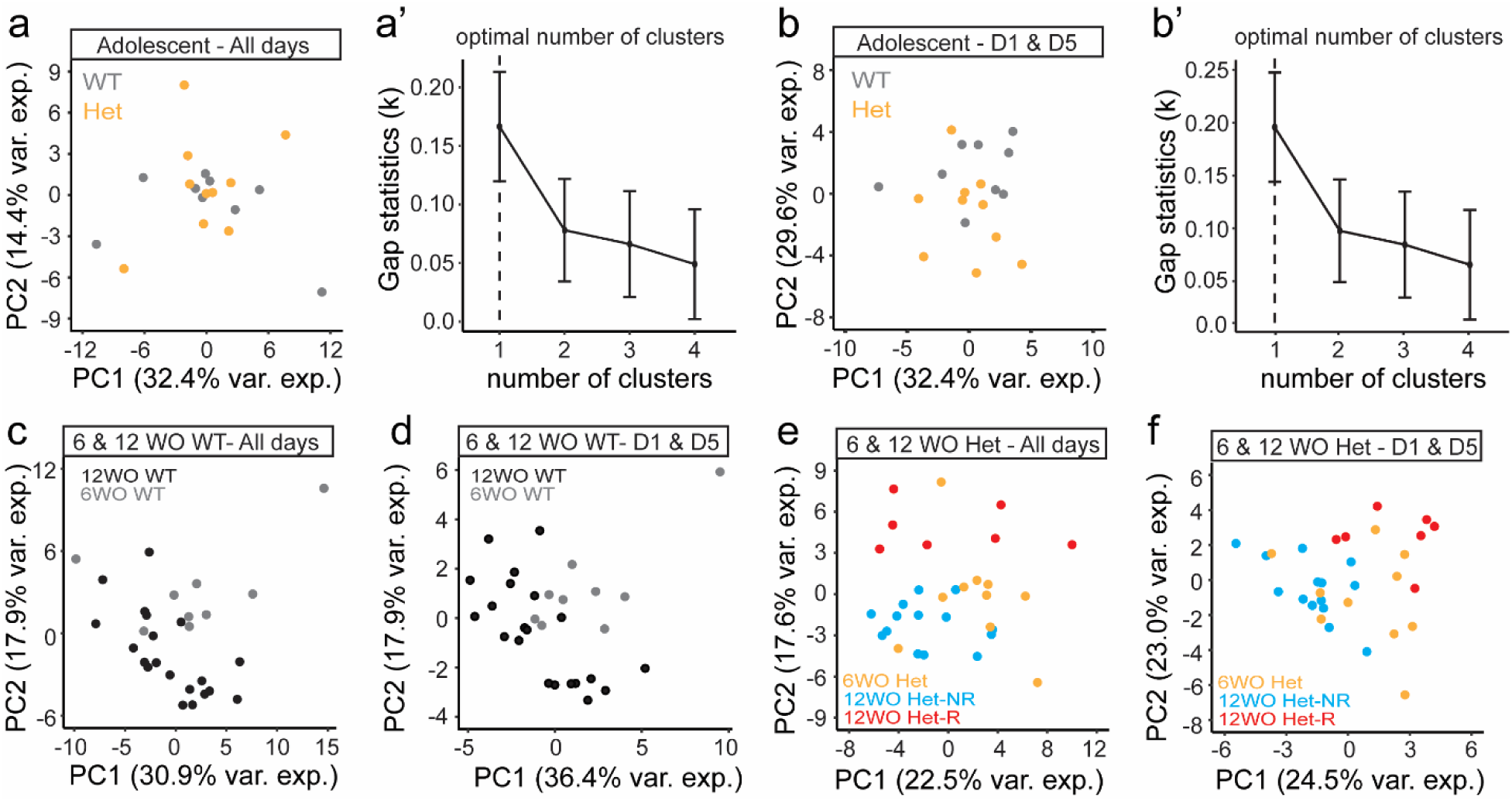
Het regression is age-specific. **a)** PCA for all days of retrieval, and **b)** only D1 & D5 of retrieval reveals no difference in PC representation between 6-Week-Old (adolescent) WT and Het [WT (9) =grey, Het (11) =orange]. **a’,b’)** Gap statistics test (see methods for details) confirms the optimal number of cluster is one (dashed line) for a’) all days of retrieval and **b’)** D1 & D5 of retrieval. c) PCA for all days of retrieval, and **d)** only D1 & D5 of retrieval reveals no difference in PC representation between adolescent WT (n=9, grey) and adult Het (n=21, black). **e)** PCA for all days of retrieval of shows adolescent Het clusters with adult Het-NR. However, D1 & D5 of retrieval **f)** shows adolescent He group with Het-NR and Het-R.

To determine if there are developmental differences in retrieval trajectories between adolescence and adulthood within each genotype, we performed PCA on all metrics for all days (Figure 6c) and D1 & D5 (Figure 6d) between adolescent and adult WT. We observed no distinct groups between adolescent WT (grey) and adult WT (black) (Figure 6c, d). We performed PCA on 13 metrics for all days (Figure 6e) and D1 & D5 (Figure 6f) between adolescent and adult Het. In all day analysis, the majority of adolescent Het grouped with adult Het-NR (Figure 6e), however, one adolescent Het grouped with adult Het-R. Interestingly, in D1 & D5 only, adolescent Het were mixed with adult Het-NR and adult Het-R (Figure 6f). Together, these results suggest possible identification of adolescent Het at risk for regression over time. Future large scale and longitudinal studies are needed to address these possibilities.

## Discussion

One of the major hurdles in diagnosing, understanding, and treating heterogeneous rare neuropsychiatric diseases is the lack of appropriate tools to robustly identify and classify complex morbidity over time and trials with individual resolution. While males are typically severely affected by MECP2 dosage, females typically exhibit less severe and more heterogeneous phenotypes due to random X-chromosome inactivation and mosaic expression of the wild-type protein. Thus, disease pathobiology caused by X-linked mutations, and by MECP2, is fundamentally different in females than males. Currently, there is minimal consensus on behavioral phenotypes in female Het mice, due to strain variations and lack of systematic analysis over different ages (Fagiolini et al., 2020; Garg et al., 2013; Krishnan et al., 2017; Mykins et al., 2023; Ribeiro & MacDonald, 2020; Samaco et al., 2013; Stearns et al., 2007; Stevenson et al., 2021). This is particularly important for designing better therapeutic interventions, as symptoms change over time in individual patients. Thus, developing and utilizing appropriate preclinical models are a necessary step. Here, we utilize pose estimation algorithm and multidimensional analysis to characterize complex social cognition phenotypes in the heterogeneous population of adolescent and adult female *Mecp2*-heterozygous mice. We found context- and age-specific regression in a subset of genotypic Het (Het-Regressors, Het-R) and an age-specific delay in efficient behavior in another subset of genotypic Het (Het-non regressors, Het-NR). These are novel results that finally achieve categorization of known, yet elusive, phenotypic heterogeneity in female X-linked disorders. Additionally, this is the first report to faithfully recapitulate regression and mild developmental delay in the apt preclinical model. This robust phenotyping in individual animals will be critical in determining the mechanisms connecting the known MECP2 mosaicism, etiology and disease progression in Rett syndrome over time, contexts and physiological states. Future work will be needed to correlate heterogeneity in disease severity to known models of MECP2 mutations, and in different relevant behavioral models (Ehrhart et al., 2021; Neul et al., 2008; Vashi & Justice, 2019). The implications of such approaches will be in establishing robust and reproducible phenotyping of female mice across labs, provide biomarkers through longitudinal progression, physiological ages and symptom severity with individual resolution, for better design for clinical interventions (Leonard et al., 2022).

### Pup retrieval task as ethologically relevant task to study complex social cognition

Maternal behavior and pup retrieval tasks are well-established paradigms to study complex dyadic interactions focusing on neurological processes such as motivation, goal-directed movements, sensory processing and perception, social communication and cognition (Alsina-Llanes et al., 2015; Beach & Jaynes, 1956; Champagne et al., 2007; Champagne & Curley, 2016; Lonstein et al., 2015; Morgan et al., 1992; Rosenblatt, 1967; Stern, 1996; Stern & Kolunie, 1989; Stevenson et al., 2021; Wiesner & Sheard, 1933). In our assay, naïve virgin female mice are co-housed with a pregnant female, 3-5 days before birth of pups. Once pups are born, we remove pups from the nest and scatter the pups in the corners of the home cage, allowing for individuals to retrieve the pups back to the nest (pup retrieval behavior), over six consecutive days. This scenario is akin to a person moving to a new place with roommates, and then motivated (or induced) to help/interact and care for newborn babies, without any prior experience, for short periods of time every day. It is a challenging task to navigate this complex social environment, with demands on cognitive flexibility, sensory processing and perception, and skilled motor sequences. The purpose of using a complex social task, which taps into a combination of innate and learned neural circuitry, in this heterogeneous population is based on the premise that brains are evolved to engage and process complex environmental and social information; thus, challenges will reveal dynamic and nuanced phenotypes in behaviors and neural processing over time and contexts, with reproducible variations in individual animals. Thus, such complexity is critical for better utilization of preclinical models of human disorders.

At the neurobiology level, much is known about the contributions of specific brain regions and neuromodulatory systems for efficient maternal care. Particularly, when nulliparous naïve female rodents are first exposed to pups, over several days, they adapt to retrieve pups, and co-housing with a pregnant female increases efficiency of pup retrieval (Cohen et al., 2011; Ehret et al., 1987; Hemel, 1973; Krishnan et al., 2017; Marlin et al., 2015; Stevenson et al., 2021). The neural circuit mechanism by which naïve female mice adapt repetitive and stereotyped motions over days to efficiently retrieve scattered pups back to the nest is still under investigation. Perturbation in dopaminergic mesolimbic pathway, ventral tegmental area and medial prefrontal cortex serotonergic neurons impair sequential pup retrieval and maternal behavior, suggesting a role for neuromodulation in motivation and reward of the pup (Gao et al., 2018, 2020; Hansen, 1994; Hansen et al., 1991a, 1991b, 1993; Keer & Stern, 1999; Wu et al., 2016; Xie et al., 2023). Dopaminergic activity increases in the ventral tegmental area and nucleus accumbens before pup contact and is required for reinforcement learning of pup retrieval over days in nulliparous female mice (Dai et al., 2022; Xie et al., 2023). Many of these established circuits have strong projections into higher cortical regions important for sensory motor integration. In this study, we have established metrics indicating naïve wild-type mice are adapting sensorimotor strategies to increase retrieval efficiency potentially to maximize the perceived social reward from pups. We hypothesize mice prioritize sensory-dependent social reward of the pups in early days of retrieval and utilize higher cortical regions to consolidate motor sequences in later days. Future work using *in-vivo* optogenetic manipulation combined with real-time pup retrieval dynamics to investigate the role of sensorimotor neural circuits is important to elucidate the underlying neural mechanisms of this relevant task.

### Neurobiology underlying sensory and social communication deficits in Rett syndrome

Longstanding work in the field of Rett syndrome has identified disruptions in cortical and subcortical regions to be major factors for sensory, motor, and social communication deficits in predominantly *Mecp2-null male* rodent models. Additionally, studies with cell-type specific deletions and “rescues” of *Mecp2* have identified particular contributions of cell-autonomous and non-cell autonomous effects in these phenotypes (Achilly, He, et al., 2021; Achilly, Wang, et al., 2021; Artoni et al., 2020; Ballinger et al., 2019; Bhattacherjee et al., 2017; Chao et al., 2010; Chapleau et al., 2009; Gemelli et al., 2006; Goffin et al., 2014; He et al., 2014; Ito-Ishida et al., 2015; Krishnan et al., 2017; Meng et al., 2016; Morello et al., 2018; Mossner et al., 2020; Orefice et al., 2016, 2019; Xu et al., 2022; Zhao et al., 2022). Specifically, mice with conditional deletion of *Mecp2* in parvalbumin+ GABAergic (PV) neurons develop a wide range of functional and behavioral RTT-like symptoms including atypical sensory-motor learning and pup retrieval behavior (Ito-Ishida et al., 2015; Krishnan et al., 2017; Mossner et al., 2020; Patrizi et al., 2020). An altered PV-network configuration interferes with experience-dependent plasticity mechanism in the brain and is associated with defective learning and plasticity (Donato et al., 2013). In the medial prefrontal cortex (mPFC) of Het, pyramidal neurons are hypo-responsive to interactions with a novel mouse or an object, and optogenetic inhibition of PV neurons rescues social deficits (Xu et al., 2022). Socially inhibited neurons in the prelimbic cortex are preferentially activated in response to a novel mouse, and this ability is impaired in MECP2-transgenic mice with overexpressed MECP2 (Zhao et al., 2022). Restoring MECP2 levels in prelimbic GABAergic interneurons rescues social novelty preference, suggesting appropriate expression of MECP2 regulates the ability to distinguish between old and novel social cues (Zhao et al., 2022). These studies collectively suggest mPFC functional hypoconnectivity in *Mecp2* mutant mice prevents discrimination between the context of social and nonsocial cues, ultimately leading to social behavioral deficits. Future work will focus on the role of mPFC in Het retrieval patterns.

While previous studies have reported regression in motor skills and respiratory function over weeks in *Mecp2* rodent models, it remains unclear when and how regression in social skills arise over time within Het (Achilly, Wang, et al., 2021; Huang et al., 2016; Veeraragavan et al., 2016). Here, using PCA and multidimensional analysis of a social cognition pup retrieval task, we observed adult Het-R are capable of adapting goal directed trajectories towards pups in early days and regress as they are unable to consolidate or maintain these efficient trajectories in later days (Figure 3). Compared to WT, Het-NR have a milder phenotype but are capable of consolidating efficient trajectories over time. Thus, Het-NR are able to compensate behaviorally, while Het-R are unable to consolidate social and sensory cues to execute efficient retrieval. Het-R regression is specific to age as we do not observe regression in adolescent Het (Mykins et al., 2023). We observe adult Het have increased interactions with novel pup stimuli, and increased interactions with textures compared to WT controls, supporting Het may have issues in cortical circuits for discriminating social and nonsocial cues (Mykins et al., 2023; Stevenson et al., 2021). Severe cortical dysregulation in these circuits may result in the inability of Het-R to consolidate changes in sensory and social cues over time or prevent reinforcement learning of the reward of the pup (Xie et al., 2023), ultimately leading to regression. *In-vivo* optogenetic manipulation in the mPFC during maternal behavior are needed to discern whether Het are unable to distinguish between socials cues of pups in later days, ultimately leading to regression. Furthermore, we have previously shown that in adult Het, pyramidal and PV neurons in the auditory cortex respond differentially to broad tones and pup calls, with lesser synaptic plasticity in the PV neurons (Lau, Krishnan, et al., 2020). The mPFC and auditory cortex reciprocally provide feedback for auditory tone discrimination, thus, optogenetic manipulation studies between these regions in the context of pup retrieval are needed to further dissect the sensory perception and motor activation pathology in Het (Rodgers & DeWeese, 2014).

Furthermore, cortical PV neurons are structurally surrounded by perineuronal nets (PNNs) which are specialized extracellular matrix structures that restrict synaptic plasticity (Carstens et al., 2016, 2021; de Vivo et al., 2013; Donato et al., 2013; Durand et al., 2012; Gogolla et al., 2009; Hou et al., 2017; Kosaka & Heizmann, 1989; Krishnan et al., 2015, 2017; Lensjø et al., 2017; Morello et al., 2018; Nakagawa et al., 1986; Patrizi et al., 2020; Pizzorusso et al., 2002; Sigal et al., 2019; Thompson et al., 2018; Ueno et al., 2017). Abnormal PNN expression and PV expression are implicated in several mouse models for neuropsychiatric disorders (Carstens et al., 2021; Emery et al., 2022; Gandhi et al., 2023; Krishnan et al., 2015, 2017; Liu et al., 2023; Pizzo et al., 2016; Reinhard et al., 2019; Sultana et al., 2021). In a CNTNAP2 mouse model for autism, degradation of excess PNNs via chondroitinase ABC (ChABC) injection in the prefrontal cortex rescues social interaction deficits (Gandhi et al., 2023). Similarly, ChABC injection in the auditory cortex of Het partially rescues pup retrieval performance, indicating value in developing therapeutics targeted at PNNs (Krishnan et al., 2017). We also reported abnormal increase in PNNs in the primary somatosensory cortex of Het which coincides with atypical tactile interactions with textures, pups, and poor pup retrieval performance (Krishnan et al., 2017; Lau, Layo, et al., 2020; Mykins et al., 2023; Stevenson et al., 2021). New tools for *in-vivo* imaging of PNNs will aid in determining how changes in PNN expression may contribute to impaired integration of social and sensory cues, and ultimately regression (Benbenishty et al., 2023).

An alternative hypothesis is Het-R may lose the ability to transfer to the next task of bringing the pups back to the nest, suggesting issues in brain regions important for motor planning and execution. In support of this hypothesis, we reported that adult Het exhibit abnormal maternal behavior task transition probabilities during pup retrieval (Stevenson et al., 2021). The locus coeruleus is important for mediating maternal behavior in mice via global release of noradrenaline between transitioning of behavioral states, suggesting dysfunction in these circuits could contribute to inefficient pup retrieval in Het. *Mecp2* mutant mice have profound physiological deficits in the locus coeruleus (Dvorkin & Shea, 2022; Howell et al., 2017; Huang et al., 2016; Taneja et al., 2009; Thomas & Palmiter, 1997). However, most of these studies focused on phenotypes in the context of respiratory dysfunction. Interestingly, DREAAD activation of mPFC pyramidal neurons ameliorate deficits in respiratory control in Het, suggesting the reciprocal feedback between locus coeruleus and mPFC may be disrupted in Het and could contribute to sensorimotor and social communication issues (Howell et al., 2017).

### Molecular and cellular mechanisms driving phenotypic variations in Het

Female patients typically survive into middle age, exhibiting sensory processing, cognitive and motor deficits throughout life (Buchanan et al., 2019; Djukic et al., 2012; Djukic & Valicenti McDermott, 2012; Key et al., 2019; LeBlanc et al., 2015; Merbler et al., 2020; Neul et al., 2010; Nomura, 2005; Peters et al., 2015; Stallworth et al., 2019; Symons et al., 2019). In these females, random X-chromosome inactivation leads to mosaic wild-type MECP2 expression in approximately 50% of the cells, while the other 50% of cells will have deficient or no MECP2 protein. The random X-chromosome inactivation patterns across body and brain, and the type of MECP2 mutations are major contributing factors to the behavioral, cellular and molecular heterogeneity in Rett syndrome. Additionally, MECP2 protein in neurons is a stable, long-life protein, with half-life of around ∼14days (Fornasiero et al., 2018; Mohar et al., 2022). Its expression increases in the postnatal brain, and correlates with the timing of critical period plasticity of sensory cortices (Braunschweig et al., 2004; Durand et al., 2012; Krishnan et al., 2015; LaSalle, 2001; Picard & Fagiolini, 2019). Recently, we reported an intriguing observation of a precocious increase in MECP2 expression within specific cell types in the primary somatosensory barrel cortex of adolescent Het that coincides with typical behavioral performance during sensory relevant tasks (Mykins et al., 2023). We speculated this may provide compensatory phenotypic benefits in the earlier age, while the inability to further increase MECP2 levels in adulthood leads to regressive phenotypes (Mykins et al., 2023). Thus, mosaic MECP2 expression or the inability to maintain cell-type specific MECP2 expression may underlie behavioral regression in our Het-R phenotype. Further in-depth analysis of cell-type specific MECP2 expression within our adult Het-NR and Het-R phenotypes is needed. Previously, we also reported lateralization of MECP2 and PNN expression in the adult primary somatosensory cortex, which could contribute to functional specialization of the adult cortex, and is dysregulated in the Het (Lau, Layo, et al., 2020). Future work will focus on identifying the nuanced expression and regulation of MECP2, an epigenetic master regulator with wide-ranging impact of gene expression.

### Application of computational neuroethology tools to understand complex endophenotypes in neuropsychiatric disorders

Basic science research using behavioral endpoints in rodent models have identified key molecular therapeutic targets for treating core symptoms of Rett syndrome patients (Garré et al., 2020; Gogliotti et al., 2016, 2017; Gualniera et al., 2021; Li et al., 2017; O’Leary et al., 2018; Smith-Hicks et al., 2017; Tai et al., 2016). Recently, the FDA approved Trofinetide as the first drug for treating disease pathology in Rett patients (Neul et al., 2022). However, the challenge of predictive power to determine which therapeutic interventions will work when and how for individual patients is the holy grail of personalized medicine. In the preclinical stages, systematic characterization of etiologically relevant behaviors in female mouse models of X-linked disorders is critical to move forward in understanding the mechanisms of complex endophenotypes in patients and identifying the efficacy of clinical therapeutics.

The emergence of computational neuroethology tools now provides a unique opportunity to shift the paradigm from single constrained reductionist behaviors to studying complex behavior in free moving animals (Shemesh & Chen, 2023). The free availability of open-source marker-less deep learning and machine learning software for behavioral segmentation using pose makes it attainable and affordable to analyze complex free moving behaviors (Berman et al., 2014; Goodwin et al., 2022; Hsu & Yttri, 2021; Lauer et al., 2022; Luxem et al., 2022; Mathis et al., 2018; Nath et al., 2019; Nilsson et al., 2020; Pereira et al., 2019, 2022; Segalin et al., 2021; Sun et al., 2021; Wiltschko et al., 2015; Winters et al., 2022). Using unsupervised machine learning, multiple groups have identified subtle, novel behavioral differences in wild-type mice, and mouse models for Alzheimer disease, stroke, and Autism, highlighting the importance of these approaches for modeling complex endophenotypes of neurological disorders in preclinical animal studies (Huang et al., 2021; Klibaite et al., 2022; Luxem et al., 2022; Segalin et al., 2021; Weber et al., 2022; Wiltschko et al., 2020).

As the field is broadening to include studies in female *Mecp2^heterozygous^* mice, there is a need for efficient methods to assess phenotypic severity. Guy et al., (2007) established a severity scoring system, however, this system is specific to rapid phenotypic onset in null males and does not apply to female Het, which have a slow and progressive phenotypic onset and display considerable heterogeneity in behaviors. Additionally, this system requires extensive monitoring over time and introduces observer bias due to its qualitative nature. We highlight the advantage of using multidimensional analysis of behavior (Figure 2) for phenotypic characterization (Figure 3) and provide a means to derive a single value output (Figure 2g) to score and evaluate Het phenotypic severity. The brevity and minimal resources for this approach make it an attractive approach to evaluate the efficacy of therapeutics in ongoing pre-clinical studies in the apt female Het mouse model. Future work will focus on clustering behavioral sequences or motifs, and manipulating neural circuitry during free moving behaviors, to determine which specific sensorimotor integration, and/or higher cognitive processes are affected during individual tasks and trials. Such studies will allow for precise testing and benchmarking of therapeutic and occupational interventions to treat neurological disorders throughout lifetime.

## Acknowledgements

We would like to thank undergraduates Ally McBryar, and Trinity Rose Schultz for technical help. We would also like to thank graduate students Logan Dunn and Jacob Elrod, and postdoc Billy Lau for technical help and intellectual discussion. This work was supported by BCMB Chancellor’s fellowships (MM), the 2021 Scholarly and Research Incentive Funds (SARIF) through the Office of Research, Innovation, & Economic Development (ORIED) at UTK (MM), BCMB James and Dora Wright Fellowship (MM), W. Sherman Kouns Excellence in Teaching award (MM), UTK 2021 Summer Undergraduate Research Group Experience Program (KK, MM, BB), UTK Rocky Top Summer Research Mentor Program (KK, MM, AJ), startup funds from the University of Tennessee—Knoxville (KK), and by the National Institute of Mental Health of the National Institutes of Health under Award Number R15MH124042 (KK).

## Contributions

All authors had full access to all the data in the study and take responsibility for the integrity of the data and accuracy of the data analysis. *Conceptualization*: KK and MM; *Methodology*: KK and MM; *Software*: MM, BB, and AJ; *Investigation*: MM, BB, and AJ; *Formal Analysis*: MM, BB, AJ *Resources*: KK, MM; Data Curation: MM, BB, AJ; Writing – *Original Draft*: MM; *Writing – Review & Editing*: KK, MM, BYBL; *Visualization*: MM; *Supervision*: KK; *Project Administration*: KK; *Funding Acquisition*: KK, MM.

## Supplemental Information

**Supplementary Figure 1.**
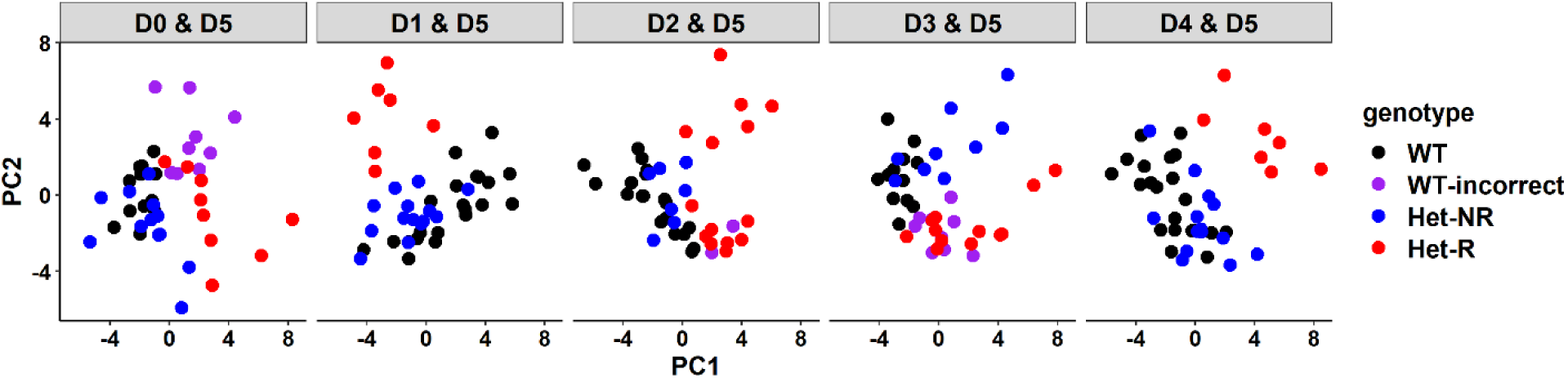
Multidimensional analysis of pairwise combinations with D5. Multidimensional analysis of pairwise combinations between (in order from left to right) D0:D5, D1:D5, D2:D5, D3:D5 and D4:D5 retrieval revealed combinations of D1 & D5 and D4 & D5 are sufficient to identify the Het-R group. Other combinations of days are insufficient because they incorrectly predict WT genotype (purple). (WT, black; WT-incorrect, purple; Het-NR, blue; Het-R, red). D0 & D5 (WT=12, WT-incorrect=9, Het-NR=12, Het-R=9), D1 & D5 (WT=21, Het-NR=14, Het-R=7), D2 & D5 (WT=19, WT-incorrect=2, Het-NR=8, Het-R=13), D3 & D5 (WT=14, WT-incorrect=7, Het-NR=10, Het-R=11), and D4 & D5 (WT=21, Het-NR=14, Het-R=7).

